# SCAPE: An AI-Driven Platform for Comprehensive Single-Cell Data Analysis

**DOI:** 10.1101/2025.11.15.688526

**Authors:** Mingke Wu, Hao Wu, Yihan Zhou, Siyuan Cheng

## Abstract

Single-cell RNA sequencing (scRNA-seq) has transformed the study of cellular heterogeneity, but downstream analysis remains fragmented and technically demanding. Current pipelines often lack analytical diversity, require programming expertise, and offer limited integration of advanced methods. To address these challenges, we developed SCAPE, an AI-driven automated and interactive platform that unifies multi-omics, multi-resource, and multi-modal single-cell data analysis. SCAPE provides platform-independent installation, customizable workflows, and integration of R- and Python-based tools. Beyond Seurat and Scanpy, it incorporates modules for transcription factor and pathway inference, pseudotime trajectory reconstruction, spatial transcriptomics deconvolution, and cell-cell communication analysis. To demonstrate SCAPE, we curated a unified atlas of lung cancer progression in human patients and mouse models, spanning primary tumors and metastatic sites. The platform enabled harmonized integration, regulatory program inference, and spatial mapping, revealing conserved epithelial programs that promote metastatic seeding and organ-specific adaptations driven by microenvironments. Collectively, SCAPE offers an accessible and comprehensive framework for single-cell analysis, providing new insights into cancer progression and broad utility across biological systems.

## Introduction

Single-cell RNA sequencing (scRNA-seq) has become an indispensable tool for dissecting cellular heterogeneity, uncovering cell states, and understanding developmental and disease processes at single-cell resolution^[1]^. For example, scRNA-seq has been widely applied in cancer heterogeneity analysis and description^[2–5]^. While upstream preprocessing pipelines, such as 10x Genomics Cell Ranger, are well-established^[6]^, the downstream analysis landscape remains complex and evolving. A variety of tools and platforms exist to process and analyze scRNA-seq data across different species, with widely used foundational packages including Seurat^[7]^ and SCANPY^[8]^ based on R and Python. These core frameworks themselves provide extensive functionality^[9]^, and building upon these foundational packages, a wide range of specialized tools have been developed to perform deeper biological interpretation. These include modules for gene ontology analysis^[10]^, transcription factor activity inference^[11]^, gene signature scoring^[12]^, pseudotime trajectory analysis^[13]^, and visualization. However, despite the richness of available tools, the scRNA-seq analysis ecosystem still faces several limitations. A major challenge lies in the lack of seamless integration across different analysis modules. Many tools function in isolation or are tightly coupled to a specific framework, making cross-platform workflows cumbersome.

Thus, a series of scRNA-seq analysis pipelines have emerged. In the early stages of scRNA-seq analysis pipeline development, many workflows were designed with the primary goal of standardizing steps. As a result, they typically focused on datasets derived from a single tissue type and implemented only a narrow range of downstream analytical strategies, often tailored to a specific biological question. For instance, the LungCellAtlas (https://github.com/LungCellAtlas/HLCA) pipeline was designed primarily to standardize the processing of lung-derived samples, with a focus on quality control and RNA velocity analysis. Similarly, eoulsan (https://github.com/GenomiqueENS/eoulsan) emphasizes robust quality control and flexible format conversion but offers limited downstream analytical capabilities. Pigx (https://github.com/BIMSBbioinfo/pigx) introduced additional features such as dimensionality reduction and basic visualization; however, the scope of these enhancements remains relatively narrow and lacks the depth required for more comprehensive single-cell analysis. Later workflows began to offer multiple solutions from raw reads to cell type annotation and basic interpretation. However, these tools often lacked depth in terms of analytical diversity and imposed a steep learning curve on users. For example, YASCP focuses on automating scRNA-seq processing but is restricted to basic tasks such as quality control, clustering, and cell annotation^[14]^. Likewise, scRNAbox (https://github.com/neurobioinfo/scrnabox) includes fundamental modules but with little room for expansion or method customization. ScRNAPip stands out by offering a wider array of downstream analyses^[15]^, including pseudotime inference, genomic instability assessment, copy number analysis, and cellular interactions inference. Nonetheless, its flexibility is significantly limited: users face substantial challenges in customizing the pipeline to suit their specific needs, and a certain level of programming expertise is required. Moreover, many of these pipelines are designed with script-based interfaces, which can be overly complex and inaccessible for users without a computational background.

To overcome the limitations of current single-cell analysis workflows including their restricted analytical diversity, poor user adaptability, and high technical barriers, we introduce an automation-driven platform that improves how single-cell data can be analyzed and interpreted. Unlike traditional pipelines that require extensive scripting or rigid, pre-defined modules, our platform features an automated inquiry system capable of dynamically guiding users through complex workflows based on diverse data inputs, parameter customization, analytical tools, and visualization options. SCAPE is an automated and interactive platform designed to unify multimodal, multi-resource, and multi-omics sequencing data analysis. Its architecture is built on two core principles: feasibility and analytical depth. The platform features an interactive setup process with modes for demonstration and custom data loading, which automatically designs tailored workflows according to the user’s specific research objectives. We have also eliminated common technical hurdles by providing a streamlined, platform-independent installation and seamlessly integrating tools from different programming environments, such as R and Python. At the same time, SCAPE dramatically expands analytical capabilities beyond conventional pipelines. It integrates a comprehensive suite of tools, from foundational frameworks like Seurat and Scanpy to cutting-edge models such as cell2sentence^[16]^ and ceLLama^[17]^ for advanced cell annotation.

Furthermore, SCAPE automates a wide range of next-generation analyses that are often siloed and complex to implement, including: inference of transcription factor and pathway activities, trajectory and pseudotime construction, cross-resource data projection, spatial transcriptomics deconvolution, and robust cell-cell communication network inference. By integrating advanced functionalities, SCAPE empowers researchers to move beyond static gene expression analysis and explore the dynamic, functional, and spatial dimensions of single-cell biology. It represents a significant step toward making powerful bioinformatics workflows more efficient, reproducible, and personalized for the entire research community.

## Results

### Establish A Single-Cell Automated Platform for Exploration Enables Interactive Analysis

SCAPE offers streamlined and platform-independent installation, including automated scripts that handle the setup of R, Python, and third-party tool dependencies across Linux, macOS, and Windows systems. This removes the need for manual environment management, reducing both setup time and error rates.

SCAPE consists of two essential components as demonstrated by Fig1 A: Interactive configuration and automated execution. The Interactive configuration offers two modes to help users get started. Demo Mode is ideal for first-time users; with a single click, it runs a complete analysis on demo dataset, showcasing the platform’s capabilities and trouble-shooting the environment setup. Custom Mode is designed for providing a user-friendly interface to specify project-specific parameters and preferences to be processed with SCAPE modules locally.

**Fig. 1.**
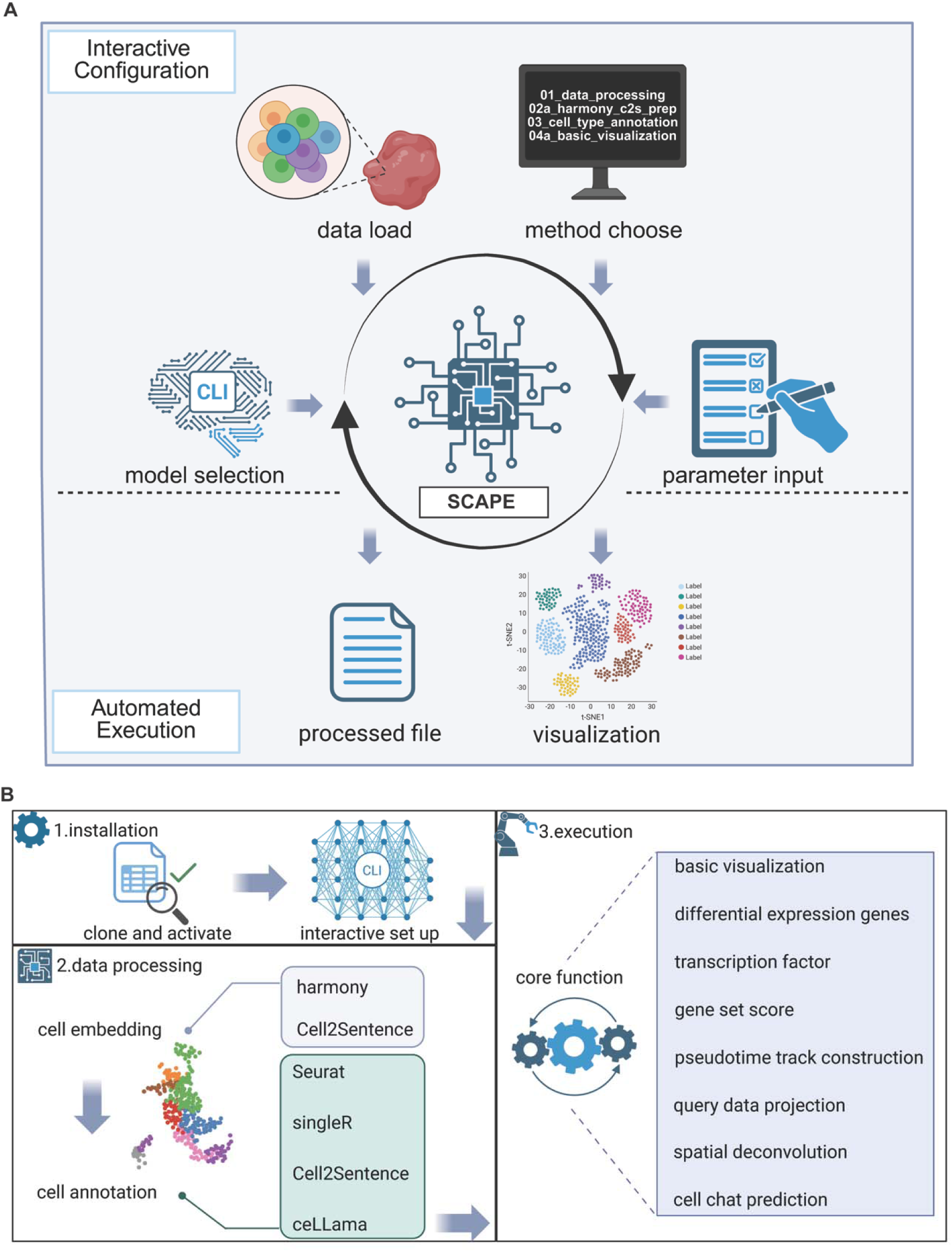
Overview of the SCAPE platform. (A) The schematic diagram illustrates the two core components and functions of. The Integrative Configuration module provides features such as data loading, SCAPE model selection, analysis method configuration, and parameter input. The Automated Execution module automatically runs the generated processing pipeline and performs visualization based on the specified parameters. (B) The flowchart demonstrates the working principle of SCAPE, mainly including installation, data processing, and execution. After cloning the repository and completing the installation, users can set various parameters through an interactive interface. The subsequent data processing, analysis, and visualization steps are then automatically executed according to the customized workflow. The complete pipeline can be found on the GitHub page.

Once the analysis plan is defined, execution is entirely managed through command-line interface (CLI) commands. The CLI enables users to control the entire workflow directly from the terminal, allowing for efficient, reproducible, and fully automated execution without the need for manual intervention. This core component automatically executes each step of the workflow according to the configured settings, enabling end-to-end automated analysis without manual intervention.

### Multi-modal Embedding and Cell Type Annotation Framework

Moving beyond the established dimensionality reduction techniques of PCA and Harmony^[18,19]^, embraces the next generation of LLM-based tools for deeper biological insight. At its core, it initiates the annotation process by leveraging the model. A key innovation is our automated execution module, which harmonizes the distinct processing environments of R and Python. This integration eliminates the technical gap between data manipulation in R and model inference in Python, ensuring a streamlined and reproducible analysis pipeline.

Besides the standard data loading, preprocessing, integration, clustering and dimensional reduction^[7]^, SCAPE offers a multi-layered approach to cell annotation, empowering users with a flexible choice of methods.

Concurrently, it employs SingleR for automated, reference-based annotation (https://github.com/dviraran/SingleR). This process assigns cell type labels by comparing the expression profile of each cell against an external, user-provided reference dataset and identifying the highest correlation scores. Besides, it also included single-cell foundation model cell2sentence to produce rich, high-dimensional cell embeddings^[16]^ to facilitate cell type annotation and cell population identification. In addition, SCAPE also offers LLM based de novo cell type annotation, powered by ceLLama^[17]^, a novel method that harnesses the power of a user-selected local model.

### SCAPE Powers Cross-Species, Multi-Omics scRNA-seq Interpretation

One of the most significant hurdles in single-cell genomics is the comparative analysis of datasets from different experiments, conditions, or species. SCAPE’s query projection module directly addresses this challenge. By implementing Seurat’s powerful reference-mapping workflow^[9]^, the pipeline can take a new query dataset and project it onto an existing, well-annotated reference. Crucially, this module includes an automated, homologene-based step for converting gene identifiers between mouse and human, a non-trivial task that is essential for cross-species comparison. This functionality enables the consistent transfer of cell type labels and facilitates the direct comparison of cellular states across species, providing an invaluable tool for translational research and the study of conserved biological processes.

A cornerstone of the SCAPE pipeline is its ability to translate gene expression into functional regulatory insights. Leveraging the decoupleR package^[20]^, our framework automates the inference of transcription factor (TF) and pathway activities from the transcriptome. Specifically, the platform integrates Unsupervised Linear Modeling (ULM) for transcription factor activity inference using the CollecTRI resource, and Multivariate Linear Modeling (MLM) for pathway activity estimation via PROGENy. To facilitate cross-species analysis and expand the platform’s utility in both human and mouse research contexts, SCAPE also includes a built-in module for automated gene ortholog conversion through homologene (https://github.com/oganm/homologene).

To further enrich the functional characterization of cell states, SCAPE incorporates a dedicated module for gene signature scoring using UCell^[12]^. This method provides robust, per-cell enrichment scores for user-defined gene sets without the confounding effects of cell-to-cell variations in dataset composition. Our pipeline enhances this analysis by allowing users to flexibly specify entire gene set collections from the MSigDB database (e.g., Hallmark H, or GO terms C5) via a single parameter. This provides an immediate, powerful readout of complex biological processes and cellular states, tailored specifically to the user’s biological questions.

Understanding dynamic cellular processes such as differentiation and activation requires moving beyond static snapshots of the transcriptome. SCAPE integrates trajectory inference through its pseudotime analysis module, which utilizes the powerful Palantir algorithm^[21]^. This analysis orders cells along a continuous trajectory, modeling their progression through a biological process.

However, after completing these analyses, generating high-quality and informative visualizations remains a major challenge for researchers. To address this, we integrated advanced visualization packages including seuratExtend, scpubr^[22,23]^, and scRNAtoolVis (https://github.com/junjunlab/scRNAtoolVis) into our pipeline. These tools enable rich and customizable outputs such as UMAP projections, gene expression feature plots, gene-level scatter plots, cell proportion barplots, volcano plots, heatmaps, and basic differential expression analyses. Importantly, users can generate these visualizations through an interactive interface by selecting custom, thereby lowering the barrier to high-quality single-cell data exploration (Fig1 B).

To integrate cellular identity with spatial organization, SCAPE incorporates the CARD algorithm for spatial transcriptomics deconvolution^[24]^. By aligning annotated scRNA-seq reference data with user-provided spatial transcriptomics matrices, CARD imputes the cell type composition at each spatial location. This integration bridges the gap between cellular heterogeneity and tissue-level architecture, enabling the reconstruction of spatially resolved cellular maps.

To elucidate intercellular communication, SCAPE leverages LIANA^[25]^, which aggregates results from multiple ligand-receptor inference methods. A distinctive feature of our implementation is its ability to execute and combine multiple algorithms in parallel, yielding consensus-based predictions that are more robust and less method-dependent. This approach enhances the reliability of cell-cell interaction inference and facilitates the identification of key signaling networks within the tissue context.

### SCAPE Reveals Transcriptomic Landscapes Across Distinct Lung Cancer Metastases

To showcase the capabilities of SCAPE, we analyzed the transcriptomic landscape of lung cancer progression across multiple metastatic stages. We first accessed data from the publicly available Human Lung Cancer Atlas, focusing on four cancer subtypes—lung adenocarcinoma, lung large-cell carcinoma, squamous cell lung carcinoma, and pleomorphic carcinoma—comprising a total of 114,000 cells^[26]^ (Fig2 A, C). In addition, we integrated single-cell RNA sequencing datasets from GEO, encompassing both human and mouse lung cancers in primary and metastatic contexts. The human datasets included primary lung tumors (lung) as well as lymph node (LN), brain, and pleural effusion (PE) metastases, totaling 208,000 cells^[27]^. Mouse datasets comprised primary tumors (lung)^[28]^ along with macrometastasis (ME), kidney, and bone, derived from models established in immunodeficient mice^[29]^, totaling 91,000 cells.

**Fig. 2.**
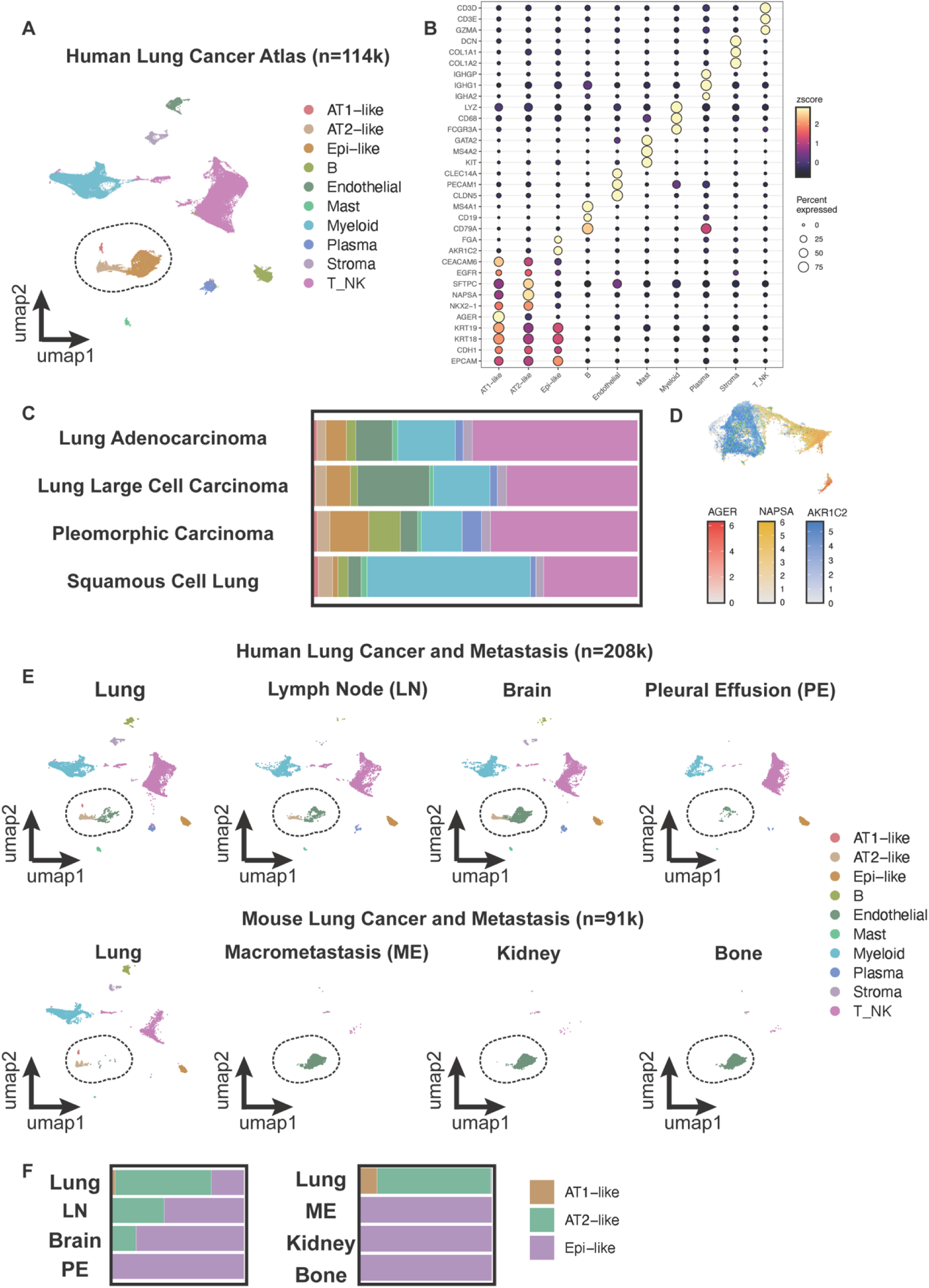
Single-cell atlas of human and mouse lung cancers. (A) UMAP visualization of the human lung cancer atlas comprising a total of 114,000 cells. (B) Dot plot shows marker genes of 10 cell types. (C) Bar plot displays the proportions of various cell types across 4 types of lung cancer. (D) Feature plot illustrates the expression of AGER, NAPSA, and AKR1C2. (E) UMAP visualization depicts metastasis landscapes of human (208,000 cells) and mouse lung cancers (91,000 cells). (F) Bar plot shows cell type proportions in different metastases of human and mouse lung cancers.

Using SCAPE, we performed comprehensive cell-type annotation of the atlas and identified ten major cellular populations (Fig2 B). Among them, epithelial lineage cells could be subdivided into AT1-like, AT2-like, and Epi-like states. All three subgroups expressed a core set of epithelial lineage markers, including KRT18, KRT19, EPCAM, and CDH1^[27,30]^. In addition to this shared program, AT1-like cells were characterized by expression of AGER^[31]^, AT2-like cells by SFTPC and NAPSA^[32]^, and Epi-like cells by FGA and AKR1C2^[33]^ (Fig2 D).

Interestingly, while all three subgroups were present in both human and mouse primary tumors, metastatic lesions exhibited a clear shift toward the Epi-like state. Brain and pleural effusion metastases in human patients, as well as all metastases in mouse models, were dominated by Epi-like tumor cells, suggesting that epithelial-like phenotypes may play a central role in metastatic colonization (Fig2 E, F).

### SCAPE Dissects Signaling Pathways and Pseudotime Dynamics of Lung Cancer

We utilized SCAPE to calculate the activity of 50 Hallmark pathways by Ucell^[34]^, and this analysis of murine and human lung cancer metastases revealed stark contrasts in metastatic (Fig3 A). In the murine model, primary tumors display a quiescent phenotype with strong suppression of proliferative pathways, whereas all metastatic lesions—regardless of organ site—exhibit uniform activation of cell cycle and MYC-related programs, suggesting a pre-programmed, proliferation-dominant metastatic program^[35]^. In human lung cancer, metastases exhibit profound heterogeneity. Pleural effusion metastases resemble the murine pattern, with extreme activation of proliferative pathways and broad suppression of other signaling programs, reflecting a proliferation-driven phenotype in a permissive, fluid-filled niche. Brain metastases, in contrast, prioritize metabolic rewiring, including oxidative phosphorylation, glycolysis, and fatty acid metabolism, over proliferation, consistent with adaptation to the high-energy, lipid-rich cerebral microenvironment. Lymph node metastases display strong immune and inflammatory signatures coupled with epithelial-mesenchymal transition and stress-response pathways, reflecting active co-evolution with host immunity^[36]^. Collectively, these findings highlight that while murine metastasis largely follows a uniform proliferative program, human metastases are shaped by organ-specific selective pressures, resulting in diverse, niche-adapted phenotypes.

**Fig. 3.**
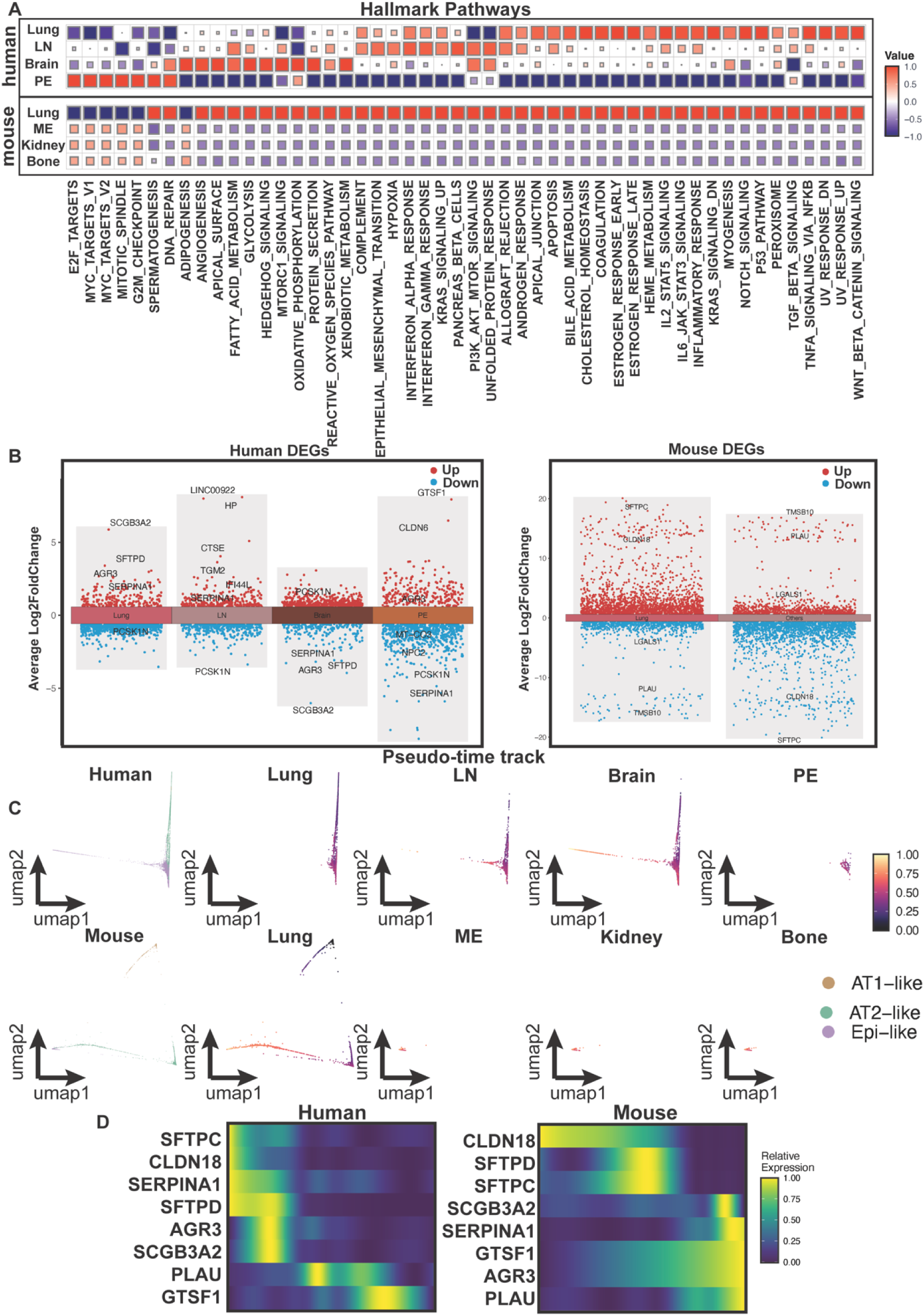
Comparative pathway and trajectory analyses of human and mouse metastatic lesions. (A) Heatmap illustrates the activity scores of human and mouse samples across 50 hallmark pathways. (B) Volcano plot presents the differentially expressed genes (DEGs) between human and mouse metastatic lesions, identifying both upregulated and downregulated genes. (C) Pseudotime UMAP visualization shows the trajectory of cells from human and mouse metastatic lesions, with pseudotime values reflecting dynamic cellular states. (D) Heatmap along the pseudotime trajectory depicts differential gene expression patterns in human and mouse.

In human lung cancer metastasis data, we systematically analyzed key gene expression changes across these metastatic sites, identifying a set of potential organ-specific metastatic markers (Fig3 B). In brain metastases, SCGB3A2 and SFTPD were significantly downregulated, suggesting that cancer cells may escape immune surveillance^[37,38]^. In lymph node metastases, TGM2 and IFI44L were markedly upregulated, reflecting enhanced proliferative and invasive capabilities, extracellular matrix remodeling, and immune evasion to promote lymphatic dissemination^[39,40]^. In pleural effusions, GTSF1 and CLDN6 were highly upregulated, potentially conferring stem-like properties and adaptation to the fluid microenvironment^[41,42]^.

In mouse metastasis data, alveolar type II marker SFTPC was strongly downregulated, indicating tumor cell dedifferentiation and acquisition of a more aggressive phenotype^[43]^(Fig3 B). CLDN18, a key tight junction protein, was also downregulated, suggesting loss of cell-cell adhesion and enhanced detachment from the primary tumor^[44]^. In contrast, LGALS1 supported angiogenesis, immune evasion, and invasiveness^[45]^, while PLAU facilitated extracellular matrix degradation, enabling tissue invasion^[46]^. These changes collectively reflect coordinated acquisition of migratory, invasive, and immune-evasive traits in metastatic tumor cells.

Using pseudotime trajectories constructed from three cell populations (AT1-like, AT2-like, and Epi-like), we set a single in situ lung cancer cell as the root (Fig3 C). In both human primary tumors, most cells exhibited low pseudotime values, indicating a state closely resembling the original tumor phenotype. In contrast, distinct clusters of cells from lymph node and brain metastases displayed markedly higher pseudotime values, suggesting divergence from the primary tumor state and acquisition of more stem-like characteristics. Notably, cells from brain metastases showed the highest pseudotime values, consistent with enhanced stemness and potential for colonization in the cerebral microenvironment. In mouse models, primary tumors exhibited heterogeneous pseudotime distributions across multiple components, whereas all three metastatic populations presented uniformly high pseudotime values, further supporting the notion that metastatic cells possess increased stem-like properties compared to their primary counterparts.

In pseudotime trajectory analysis (Fig 3D), early-expressed genes in human lung cancer, including SCGB3A2, SFTPD, and CLDN18, are primarily associated with alveolar epithelial differentiation and cell adhesion, suggesting that tumor cells retain partial epithelial characteristics in the early stages^[37,38,44]^. In contrast, late-expressed genes such as PLAU and GTSF1 are linked to extracellular matrix remodeling and stem-like features, indicating that tumor cells progressively acquire migratory, invasive, and immune-evasive capabilities during progression and metastasis^[42]^ ^[46]^. Similarly, in the mouse lung cancer model, early expression of CLDN18 and SFTPD reflects the retention of epithelial traits^[37,44]^, whereas late-expressed genes are involved in cytoskeletal remodeling, invasiveness, and adaptation to the metastatic niche^[42,46]^. Overall, these pseudotime dynamics suggest a stage-specific transition in metastatic lung cancer cells from epithelial-like states toward migratory, invasive, and microenvironment-adapted phenotypes.

### SCAPE Enables the Inference of Cellular Communication and Transcription Factor Activity

We applied SCAPE to predict cell–cell interactions using five complementary methods implemented in LIANA (natmi, connectome, logfc, sca, and cellphonedb). Cellular communication analysis using revealed distinct, metastasis-specific ligand–receptor networks in human and mouse lung cancer (Fig4 A, C). In lymph node metastases, AT2-like cells engaged in a broad expansion of interactions, such as TNC–SDC1/4 and TIMP1–CD63, which are associated with extracellular matrix remodeling, cell adhesion, and metastatic niche conditioning^[47,48]^. These interactions may facilitate immune evasion and promote metastatic colonization in the immunologically active lymph node microenvironment. In brain metastases, classical matrix–receptor interactions including COL1A1–DDR2 and COL6A2–SDC1 were markedly reduced, suggesting a loss of canonical stromal anchoring and a shift toward alternative, metabolically driven adaptation strategies within the neural niche^[49,50]^. By contrast, pleural effusion metastases exhibited striking reinforcement of AT2-like autocrine signaling, exemplified by CEACAM–EGFR and NDP–FZD4 interactions, pointing to a self-sustaining proliferative program in the relatively permissive fluid microenvironment^[51,52]^(Fig4 B). In murine metastases, however, interaction patterns were highly conserved across sites, dominated by AT2-like signals such as CDH1–ERBB3 and CEACAM6–EGFR, mirroring the uniform proliferative program revealed by hallmark analysis^[53,54]^(Fig4 D). Together, these findings highlight that while murine metastases are governed by a stereotyped, proliferation-centered program, human metastases undergo niche-specific rewiring of cell–cell communication that underlies their phenotypic diversity.

**Fig. 4.**
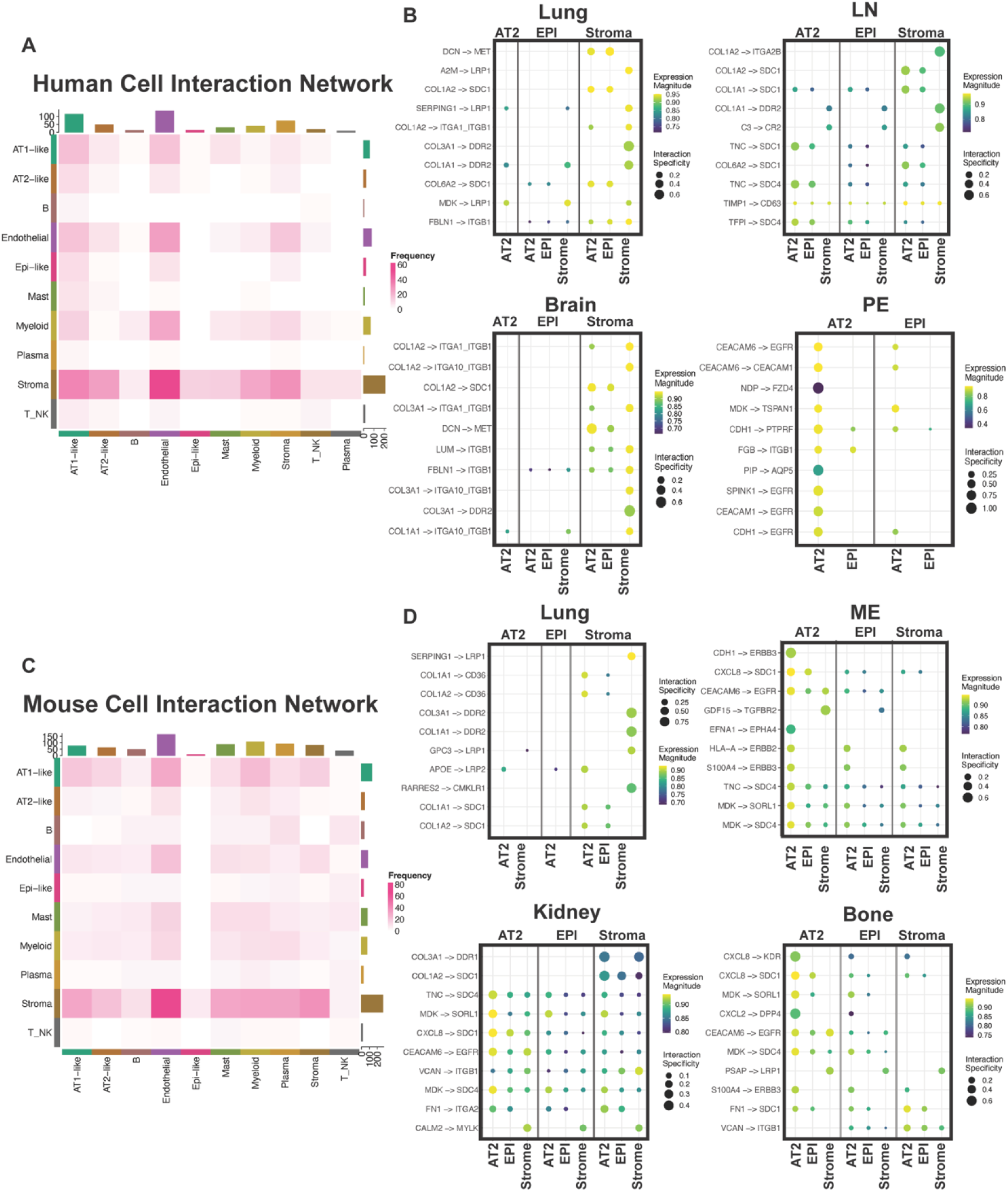
Cell–cell communication analysis in human and mouse metastatic lesions. (A) Heatmap illustrates the interaction strengths among different human cell types. (B) Dot plot shows the specific ligand–receptor interaction strengths across different human metastatic lesions. (C) Heatmap illustrates the interaction strengths among different mouse cell types. (D) Dot plot shows the specific ligand–receptor interaction strengths across different mouse metastatic lesions.

SCAPE further allowed us to infer the top transcription factors shaping regulatory states across distinct metastatic sites in lung cancer, revealing strikingly divergent transcriptional programs driven by microenvironmental contexts (Fig5 A). In human pleural effusions, tumor cells displayed a unique activation of the TGF-β/BMP axis, with exceptionally high SMAD9 and SMAD5, consistent with a survival program tailored to the inflammatory, cytokine-rich pleural cavity^[55]^. By contrast, lymph node metastases were defined by a metastasis–immune evasion module involving ZBTB38, ZNF217, and CTBP2, enabling tumor cells to simultaneously enhance intrinsic invasive potential and evade the highly immunocompetent nodal milieu^[56–58]^. Primary tumors maintained a lineage-survival transcriptional core centered on NKX2-1 and TTF1, as well as cooperating partners such as ETV5 and FOXM1, reflecting lineage addiction–driven proliferation. Strikingly, brain metastases exhibited a largely transcriptionally silent landscape, lacking strong TF activation and instead suggesting alternative adaptive strategies, such as post-transcriptional regulation or metabolic rewiring. Comparative analysis in a murine lung cancer model further revealed an amplified activation of oncogenic drivers including PLAGL2 and NKX2-1 in primary tumors^[59,60]^, coupled with coordinated suppression of differentiation-associated (SOX9, DACH1)^[61,62]^ and EMT-associated (ZEB1, LEF1)^[63,64]^ TFs in murine metastatic sites, thereby highlighting that in murine model, primary tumors exist in a highly proliferative epithelial state in which EMT may not be required for subsequent metastatic dissemination. Moreover, such a rigid epithelial program likely constrains cellular plasticity, thereby limiting the ability of tumor cells to adapt to distant metastatic microenvironments.

**Fig. 5.**
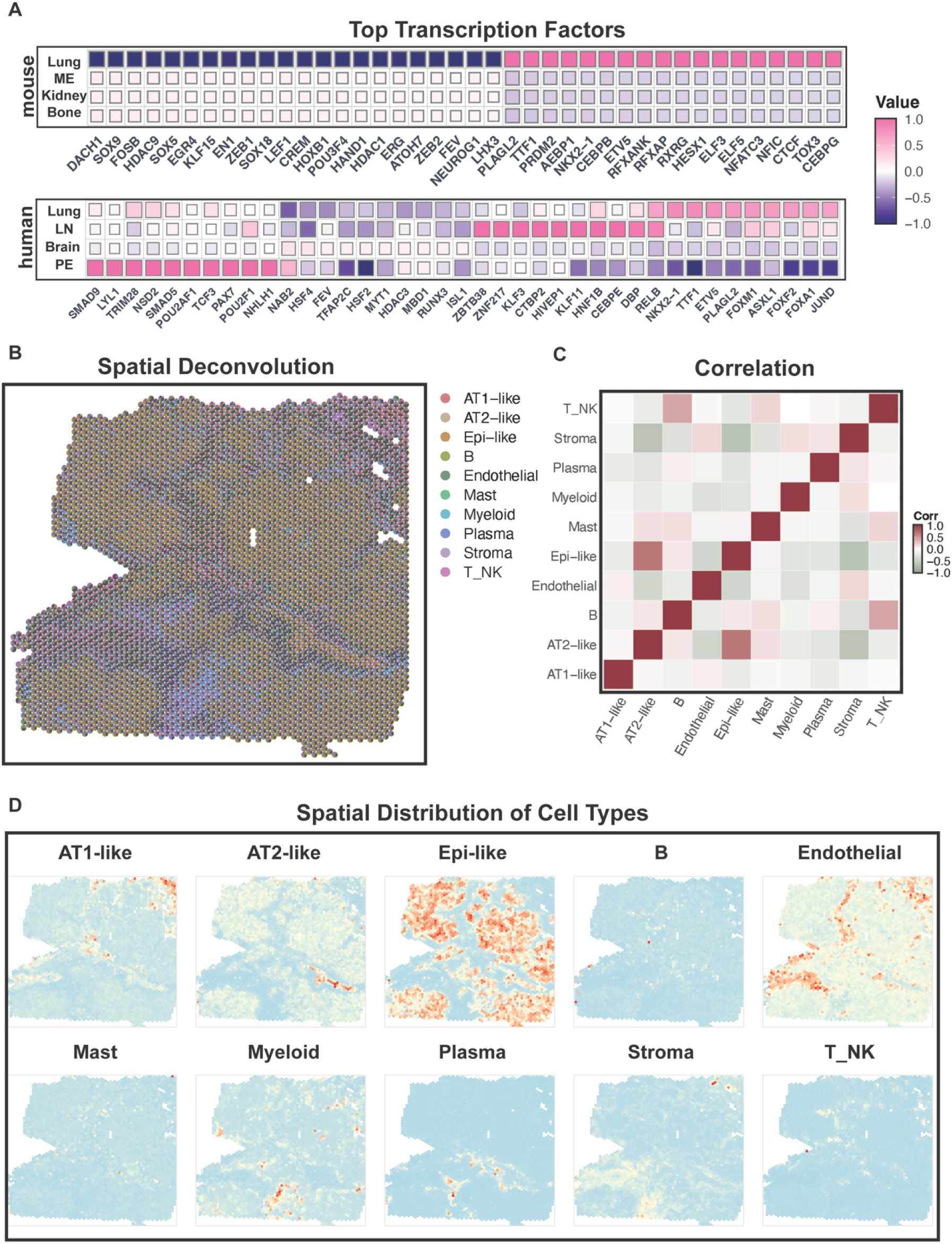
Transcriptional regulation and spatial organization in human and mouse samples. (A) Heatmap illustrates the activity of different transcription factors in human and mouse samples. (B) Spatial pie plots show the cellular composition within each spatial spot. (C) Correlation heatmap depicts the interaction strength between each pair of cell types. (D) Spatial transcriptomic map displays the spatial distribution of 10 cell types across the tissue.

### Integration of Single-Cell and Spatial Transcriptomic Data Facilitated by SCAPE

We downloaded and processed spatial transcriptomic data of Lung Cancer (https://www.10xgenomics.com/datasets/human-lung-cancer-11-mm-capture-area-ffpe-2-standar d) from the 10x Genomics database. Deconvolution was performed using CARD, with reference single-cell data derived from our previous project. The mapping results demonstrated high fidelity, as the vast majority of spots were confidently identified as tumor cells, reflecting the accuracy of our deconvolution approach (Fig5 B). Correlation analysis further revealed strong intra-group interactions within each metastatic site, providing complementary insights beyond those predicted by cell interaction prediction. Notably, the robust interactions between Epi-like and AT2-like cells were consistently observed, in agreement with previous findings (Fig5 C).

Upon examining the spatial distribution of non-malignant cell types, we observed that plasma cells and T_NK (T/NK) cells were interspersed among the tumor cells. This intermingling suggests active immune infiltration within the tumor microenvironment, potentially indicating ongoing antitumor immune responses, local recognition of tumor antigens, or dynamic tumor-immune interactions^[65,66]^. We also noted a relatively high abundance of myeloid cells, which showed partial spatial overlap with plasma cells. This co-localization may point to immune crosstalk between innate and adaptive compartments, possibly reflecting immunoregulatory or immunosuppressive niches within the tumor microenvironment^[67]^. Myeloid cells, depending on their polarization state, can either support antitumor immunity or promote tumor progression through suppression of T/NK cell function^[68,69]^ (Fig5 D). The spatial proximity of plasma and myeloid cells may therefore highlight microdomains where immune modulation is actively occurring, which could have implications for understanding tumor progression and designing targeted therapies.

## Discussion

In this study, we introduce SCAPE, a next-generation analytical platform tailored for comprehensive and automated single-cell multi-omics analysis. Through seamless integration of state-of-the-art bioinformatics tools, SCAPE demonstrates substantial improvements in usability, analytical flexibility, and biological interpretability.

One of the most significant contributions of SCAPE is its intelligent, interactive, and code-free workflow, which enables researchers with varying computational backgrounds to engage in high-level single-cell analysis. Unlike traditional pipelines that require extensive scripting and tool-specific knowledge, SCAPE employs a prompt-driven interface and smart parameter tuning, substantially lowering the barrier to entry while preserving analytical depth and reproducibility. Our integration of LLM-based annotation tools such as cell2sentence and ceLLama offers a powerful complement to conventional reference-based methods like SingleR. While reference-dependent strategies provide reliable annotations in well-characterized systems, they often falter when encountering novel or heterogeneous cell states. LLM-based models, in contrast, enable context-aware, reference-free predictions that are especially valuable in immuno-oncology and developmental biology, where cellular states may be highly dynamic or previously uncharacterized.

Moreover, the ability of SCAPE to span across modalities, platforms, and species marks a major step toward unified multi-omics analysis. The inclusion of trajectory inference, transcription factor and pathway activity estimation, and gene signature scoring enhances the platform’s capacity to dissect dynamic biological processes such as differentiation and activation. The automated homologene-based gene ortholog conversion, along with the Seurat reference projection module, expands this utility to cross-species and cross-condition comparisons—an increasingly essential capability in translational and evolutionary biology.

Importantly, we validated the full approach to construct a comprehensive atlas of lung cancer progression across primary and metastatic contexts in both humans and mice, revealing both conserved programs and organ-specific adaptations. A key finding was the marked divergence between human and murine metastases: while mouse models exhibited a uniform, proliferation-dominant program across all metastatic sites, human metastases displayed striking heterogeneity shaped by local microenvironments. This distinction likely reflects the absence of adaptive immune pressure in immunodeficient murine models, in contrast to the complex ecological constraints of human disease. Across species, we observed a consistent enrichment of Epi-like tumor states in metastases, suggesting that epithelial phenotypes confer advantages for colonization, although in mice this rigid epithelial program was coupled with suppression of EMT-associated regulators, thereby limiting plasticity, whereas human metastases retained greater flexibility and rewiring potential.

Despite these advancements, some limitations remain. While the platform supports multiple modalities, integration of additional data types such as scATAC-seq and proteomics could further enhance its scope. Besides, while the command-line interface is intuitive, future versions of could benefit from a full graphical user interface (GUI) to further democratize access.

In conclusion, SCAPE presents a powerful, scalable, and user-friendly solution for modern single-cell analysis. By tightly coupling an automated pipeline with cutting-edge analytical tools, SCAPE not only accelerates the pace of discovery but also reshapes how computational biology can be practiced in the era of automation and advanced computational tools. In addition, this work demonstrates the power of SCAPE to resolve the cellular and molecular landscapes of lung cancer metastasis, revealing both conserved epithelial programs and organ-specific adaptations that shape metastatic trajectories and pointing to potential therapeutic opportunities tailored to distinct metastatic niches.

## Methods

### SCAPE Platform Overview

The pipeline in this study was executed in an automated manner via the command line, using Git Bash on Windows, Terminal on macOS, and standard shell environments on Linux. The workflow was initiated by providing a parameter file to a Python script, enabling consistent and reproducible analysis across different operating systems. The pipeline was primarily developed using R (version 4.2.3) and Python (version 3.9). The Python package cell2sentence (version 1.1.0) was utilized in the analysis. A full list of R package dependencies and their respective versions is available in the supplementary file r_package_versions.csv.

### Quality Control and Preprocessing

Standard quality control was applied, and users could optionally exclude cell cycle-related cells and doublets using Seurat and scDblFinder (https://github.com/plger/scDblFinder) packages.

### Cell Embeddings and Annotation

The process begins with the generation of high-dimensional cell embeddings using either a straightforward PCA-Harmony pipeline or the cell2sentence (https://github.com/vandijklab/cell2sentence) model. Unsupervised clustering is then performed using the Seurat package (https://github.com/satijalab/seurat), based on Harmony-corrected principal components, to define initial cell groupings. For reference-based annotation, SingleR (https://github.com/dviraran/SingleR) is employed to compare expression profiles against user-provided reference datasets, such as the Azimuth PBMC reference. To complement this, cell2sentence is further utilized for large LLM-based cell type prediction, offering an independent, model-driven annotation that does not rely on external references-particularly valuable for identifying novel or heterogeneous cell populations. For advanced de novo annotation, ceLLama (https://github.com/CelVoxes/ceLLama) is integrated into the pipeline to generate descriptive cell type labels by interpreting the combined biological functions of marker genes, leveraging a user-selected local LLM.

### Advanced Visualizations

To enhance and customize visualizations beyond base Seurat plotting functions, advanced packages like SeuratExtend (https://github.com/huayc09/SeuratExtend), SCpubr (https://github.com/enblacar/SCpubr), and scRNAtoolVis (https://github.com/junjunlab/scRNAtoolVis) are integrated. These enable rich outputs such as UMAP projections, gene expression feature plots, gene-level scatter plots, cell proportion barplots, volcano plots, and heatmaps. Users can generate these visualizations through an interactive interface by selecting custom options or using an AI-assisted agent with natural language input.

### Transcription Factor and Pathway Activity Inference

The decoupleR (https://saezlab.github.io/decoupleR/) package is leveraged to automate the inference of transcription factor (TF) and pathway activities. Specifically, Unsupervised Linear Modeling (ULM) with the CollecTRI (https://github.com/saezlab/CollecTRI) resource is used for TF activity inference, and Multivariate Linear Modeling (MLM) via PROGENy (https://github.com/saezlab/progeny) is used for pathway activity estimation.

### Cross-Species Transformation

A built-in module for automated gene ortholog conversion through homologene (https://github.com/oganm/homologene) is included to facilitate cross-species analysis in human and mouse research.

### Gene Signature Scoring

The UCell (https://github.com/carmonalab/UCell) method is incorporated for robust, per-cell enrichment scores for user-defined gene sets, allowing flexible specification of gene set collections from the MSigDB (https://www.gsea-msigdb.org/gsea/msigdb) database (e.g., Hallmark H, GO terms C5).

### Pseudotime Trajectory Analysis

Trajectory inference is performed through a pseudotime analysis module, utilizing the Palantir (https://github.com/dpeerlab/Palantir) algorithm to order cells along a continuous trajectory, modeling their progression through biological processes.

### Cross-Resource Data Projection

The query projection module implements Seurat’s reference-mapping workflow to project new query datasets onto existing, well-annotated references (https://github.com/satijalab/seurat).

### Spatial Transcriptomics Deconvolution

The CARD (https://github.com/YMa-lab/CARD) algorithm is incorporated to integrate cellular identity with spatial organization. It aligns annotated scRNA-seq reference data with user-provided spatial transcriptomics matrices to impute cell type composition at each spatial location.

### Cell-Cell Communication Network Inference

LIANA (https://github.com/saezlab/liana) is leveraged for intercellular communication analysis, aggregating results from multiple ligand-receptor inference methods. This implementation executes and combines multiple algorithms in parallel for consensus-based predictions, enhancing robustness and reliability.

## Code Availability

All code used in this study is available and openly accessible at the following GitHub repository: https://github.com/sorrymaker03/SCAPE.

## Data Availability

To demonstrate and validate our analytical pipeline, we downloaded single-cell RNA-seq datasets from public databases, including GSE131907^[70]^, GSE123903^[71]^, and GSE222901^[72]^. From GSE131907, we selected primary tumors and three metastatic sites: lymph node, brain, and pleural effusion. From GSE123903, we included three metastatic sites: macrometastases in the kidney and bone. From GSE222901, we selected 3 primary lung cancer samples, consistent with the previous mouse metastatic data. Additionally, we used all cancer-related data from the Human Cell Atlas as reference^[73]^, including lung adenocarcinoma, lung large-cell carcinoma, squamous cell lung carcinoma, and pleomorphic carcinoma (https://data.humancellatlas.org/hca-bio-networks/lung/atlases/lung-v1-0). In addition, an annotated scRNA-seq dataset was used as a reference to deconvolve a spatial transcriptomics dataset derived from a human lung cancer sample (Neuroendocrine Carcinoma), provided by 10x Genomics (https://www.10xgenomics.com/datasets/human-lung-cancer-11-mm-capture-area-ffpe-2-standar d).

## CRediT Author Statement

**Mingke Wu**: Conceptualization, Software, Data curation, Visualization, Writing-Original draft preparation; **Hao Wu**: Visualization, Writing-Original draft preparation; **Yihan Zhou**: Writing-Reviewing and Editing; **Siyuan Cheng**: Conceptualization, Methodology, Writing-Reviewing and Editing.

## Reference

1. Tang, F., Barbacioru, C., Wang, Y., Nordman, E., Lee, C., Xu, N., Wang, X., Bodeau, J., Tuch, B. B., Siddiqui, A., Lao, K., & Surani, M. A. (2009). mRNA-Seq whole-transcriptome analysis of a single cell. Nature Methods, 6(5), 377–382. 10.1038/nmeth.1315

2. Xu, Y., Yang, Y., Wang, Z., Sjöström, M., Jiang, Y., Tang, Y., Cheng, S., Deng, S., Wang, C., Gonzalez, J., Johnson, N. A., Li, X., Li, X., Metang, L. A., Mukherji, A., Xu, Q., Tirado, C. R., Wainwright, G., Yu, X., … Mu, P. (2024). ZNF397 Deficiency Triggers TET2-Driven Lineage Plasticity and AR-Targeted Therapy Resistance in Prostate Cancer. Cancer Discovery, 14(8), 1496–1521. 10.1158/2159-8290.CD-23-0539

3. Jiang, Y., Cheng, S., Li, L., Fraidenburg, M., Kim, I. Y., Deng, S., & Mu, P. (2025). Notch as a Driver of Lineage Plasticity and Therapeutic Target in Enzalutamide-Resistant Prostate Cancer. Cancer Biology. 10.1101/2025.05.26.656166

4. Deng, S., Wang, C., Wang, Y., Xu, Y., Li, X., Johnson, N. A., Mukherji, A., Lo, U.-G., Xu, L., Gonzalez, J., Metang, L. A., Ye, J., Tirado, C. R., Rodarte, K., Zhou, Y., Xie, Z., Arana, C., Annamalai, V., Liu, X., … Mu, P. (2022). Ectopic JAK–STAT activation enables the transition to a stem-like and multilineage state conferring AR-targeted therapy resistance. Nature Cancer, 3(9), 1071–1087. 10.1038/s43018-022-00431-9

5. Li, X., Wang, Y., Deng, S., Zhu, G., Wang, C., Johnson, N. A., Zhang, Z., Tirado, C. R., Xu, Y., Metang, L., Gonzalez, J., Mukherji, A., Ye, J., Yang, Y., Peng, W., Tang, Y., Hofstad, M., Xie, Z., Yoon, H., … Mu, P. (2023). Loss of SYNCRIP unleashes APOBEC-driven mutagenesis, tumor heterogeneity, and AR-targeted therapy resistance in prostate cancer. Cancer Cell, 41(8), 1427–1449.e12. 10.1016/j.ccell.2023.06.010

6. Zheng, G. X. Y., Terry, J. M., Belgrader, P., Ryvkin, P., Bent, Z. W., Wilson, R., Ziraldo, S. B., Wheeler, T. D., McDermott, G. P., Zhu, J., Gregory, M. T., Shuga, J., Montesclaros, L., Underwood, J. G., Masquelier, D. A., Nishimura, S. Y., Schnall-Levin, M., Wyatt, P. W., Hindson, C. M., … Bielas, J. H. (2017). Massively parallel digital transcriptional profiling of single cells. Nature Communications, 8(1), 14049. 10.1038/ncomms14049

7. Satija, R., Farrell, J. A., Gennert, D., Schier, A. F., & Regev, A. (2015). Spatial reconstruction of single-cell gene expression data. Nature Biotechnology, 33(5), 495–502. 10.1038/nbt.3192

8. Wolf, F. A., Angerer, P., & Theis, F. J. (2018). SCANPY: Large-scale single-cell gene expression data analysis. Genome Biology, 19(1), 15. 10.1186/s13059-017-1382-0

9. Hao, Y., Stuart, T., Kowalski, M. H., Choudhary, S., Hoffman, P., Hartman, A., Srivastava, A., Molla, G., Madad, S., Fernandez-Granda, C., & Satija, R. (2024). Dictionary learning for integrative, multimodal and scalable single-cell analysis. Nature Biotechnology, 42(2), 293–304. 10.1038/s41587-023-01767-y

10. Huang, D. W., Sherman, B. T., Tan, Q., Kir, J., Liu, D., Bryant, D., Guo, Y., Stephens, R., Baseler, M. W., Lane, H. C., & Lempicki, R. A. (2007). DAVID Bioinformatics Resources: Expanded annotation database and novel algorithms to better extract biology from large gene lists. Nucleic Acids Research, 35(suppl_2), W169–W175. 10.1093/nar/gkm415

11. Badia-i-Mompel, P., Vélez Santiago, J., Braunger, J., Geiss, C., Dimitrov, D., Müller-Dott, S., Taus, P., Dugourd, A., Holland, C. H., Ramirez Flores, R. O., & Saez-Rodriguez, J. (2022). decoupleR: Ensemble of computational methods to infer biological activities from omics data. Bioinformatics Advances, 2(1), vbac016. 10.1093/bioadv/vbac016

12. Andreatta, M., & Carmona, S. J. (2021). UCell: Robust and scalable single-cell gene signature scoring. Computational and Structural Biotechnology Journal, 19, 3796–3798. 10.1016/j.csbj.2021.06.043

13. Qiu, X., Hill, A., Packer, J., Lin, D., Ma, Y.-A., & Trapnell, C. (2017). Single-cell mRNA quantification and differential analysis with Census. Nature Methods, 14(3), 309–315. 10.1038/nmeth.4150

14. Felipe Marques de Almeida, Alexander Peltzer, Gregor Sturm, heylf, Olga Botvinnik, nf-core bot, Dongze He, Nico Trummer, Kevin Menden, Adam Talbot, gennadyFauna, Sangram K Sahu, Gisela Gabernet, Robert Syme, Pol Avec, Maxime U Garcia, Harshil Patel, Tom Kelly, Peter Bailey, … Wei-An Chen. (2024). nf-core/scrnaseq: 2.6.0 (Version 2.6.0) [Computer software]. Zenodo. 10.5281/ZENODO.3568187

15. Xu, L., Zhang, J., He, Y., Yang, Q., Mu, T., Guo, Q., Li, Y., Tong, T., Chen, S., & Ye, R. D. (2023). ScRNAPip: A systematic and dynamic pipeline for single-cell RNA sequencing analysis. iMeta, 2(4), e132. 10.1002/imt2.132

16. Levine, D., Rizvi, S. A., Lévy, S., Pallikkavaliyaveetil, N., Zhang, D., Chen, X., Ghadermarzi, S., Wu, R., Zheng, Z., Vrkic, I., Zhong, A., Raskin, D., Han, I., De Oliveira Fonseca, A. H., Caro, J. O., Karbasi, A., Dhodapkar, R. M., & Van Dijk, D. (2023). Cell2Sentence: Teaching Large Language Models the Language of Biology. Bioinformatics. 10.1101/2023.09.11.557287

17. Choi, H., Park, J., Kim, S., Kim, J., Lee, D., Bae, S., Shin, H., & Lee, D. (2024). CELLama: Foundation Model for Single Cell and Spatial Transcriptomics by Cell Embedding Leveraging Language Model Abilities. Bioinformatics. 10.1101/2024.05.08.593094

18. Korsunsky, I., Millard, N., Fan, J., Slowikowski, K., Zhang, F., Wei, K., Baglaenko, Y., Brenner, M., Loh, P., & Raychaudhuri, S. (2019). Fast, sensitive and accurate integration of single-cell data with Harmony. Nature Methods, 16(12), 1289–1296. 10.1038/s41592-019-0619-0

19. Jolliffe, I. T., & Cadima, J. (2016). Principal component analysis: A review and recent developments. Philosophical Transactions. Series A, Mathematical, Physical, and Engineering Sciences, 374(2065), 20150202. 10.1098/rsta.2015.0202

20. Badia-i-Mompel, P., Vélez Santiago, J., Braunger, J., Geiss, C., Dimitrov, D., Müller-Dott, S., Taus, P., Dugourd, A., Holland, C. H., Ramirez Flores, R. O., & Saez-Rodriguez, J. (2022). decoupleR: Ensemble of computational methods to infer biological activities from omics data. Bioinformatics Advances, 2(1), vbac016. 10.1093/bioadv/vbac016

21. Setty, M., Kiseliovas, V., Levine, J., Gayoso, A., Mazutis, L., & Pe’er, D. (2019). Characterization of cell fate probabilities in single-cell data with Palantir. Nature Biotechnology, 37(4), 451–460. 10.1038/s41587-019-0068-4

22. Blanco-Carmona, E. (2022). Generating publication ready visualizations for Single Cell transcriptomics using SCpubr. Bioinformatics. 10.1101/2022.02.28.482303

23. Hua, Y., Weng, L., Zhao, F., & Rambow, F. (2024). SeuratExtend: Streamlining Single-Cell RNA-Seq Analysis Through an Integrated and Intuitive Framework. Bioinformatics. 10.1101/2024.08.01.606144

24. Ma, Y., & Zhou, X. (2022). Spatially informed cell-type deconvolution for spatial transcriptomics. Nature Biotechnology, 40(9), 1349–1359. 10.1038/s41587-022-01273-7

25. Dimitrov, D., Türei, D., Garrido-Rodriguez, M., Burmedi, P. L., Nagai, J. S., Boys, C., Ramirez Flores, R. O., Kim, H., Szalai, B., Costa, I. G., Valdeolivas, A., Dugourd, A., & Saez-Rodriguez, J. (2022). Comparison of methods and resources for cell-cell communication inference from single-cell RNA-Seq data. Nature Communications, 13(1), 3224. 10.1038/s41467-022-30755-0

26. Sikkema, L., Ramírez-Suástegui, C., Strobl, D. C., Gillett, T. E., Zappia, L., Madissoon, E., Markov, N. S., Zaragosi, L.-E., Ji, Y., Ansari, M., Arguel, M.-J., Apperloo, L., Banchero, M., Bécavin, C., Berg, M., Chichelnitskiy, E., Chung, M., Collin, A., Gay, A. C. A., … Theis, F. J. (2023). An integrated cell atlas of the lung in health and disease. Nature Medicine, 29(6), 1563–1577. 10.1038/s41591-023-02327-2

27. Kim, N., Kim, H. K., Lee, K., Hong, Y., Cho, J. H., Choi, J. W., Lee, J.-I., Suh, Y.-L., Ku, B. M., Eum, H. H., Choi, S., Choi, Y.-L., Joung, J.-G., Park, W.-Y., Jung, H. A., Sun, J.-M., Lee, S.-H., Ahn, J. S., Park, K., … Lee, H.-O. (2020). Single-cell RNA sequencing demonstrates the molecular and cellular reprogramming of metastatic lung adenocarcinoma. Nature Communications, 11(1), 2285. 10.1038/s41467-020-16164-1

28. Han, G., Sinjab, A., Rahal, Z., Lynch, A. M., Treekitkarnmongkol, W., Liu, Y., Serrano, A. G., Feng, J., Liang, K., Khan, K., Lu, W., Hernandez, S. D., Liu, Y., Cao, X., Dai, E., Pei, G., Hu, J., Abaya, C., Gomez-Bolanos, L. I., … Kadara, H. (2024). An atlas of epithelial cell states and plasticity in lung adenocarcinoma. Nature, 627(8004), 656–663. 10.1038/s41586-024-07113-9

29. Laughney, A. M., Hu, J., Campbell, N. R., Bakhoum, S. F., Setty, M., Lavallée, V.-P., Xie, Y., Masilionis, I., Carr, A. J., Kottapalli, S., Allaj, V., Mattar, M., Rekhtman, N., Xavier, J. B., Mazutis, L., Poirier, J. T., Rudin, C. M., Pe’er, D., & Massagué, J. (2020). Regenerative lineages and immune-mediated pruning in lung cancer metastasis. Nature Medicine, 26(2), 259–269. 10.1038/s41591-019-0750-6

30. Wu, F., Fan, J., He, Y., Xiong, A., Yu, J., Li, Y., Zhang, Y., Zhao, W., Zhou, F., Li, W., Zhang, J., Zhang, X., Qiao, M., Gao, G., Chen, S., Chen, X., Li, X., Hou, L., Wu, C., … Zhou, C. (2021). Single-cell profiling of tumor heterogeneity and the microenvironment in advanced non-small cell lung cancer. Nature Communications, 12(1), 2540. 10.1038/s41467-021-22801-0

31. Kanagaki, S., Ikeo, S., Suezawa, T., Yamamoto, Y., Seki, M., Hirai, T., Hagiwara, M., Suzuki, Y., & Gotoh, S. (2021). Directed induction of alveolar type I cells derived from pluripotent stem cells via Wnt signaling inhibition. Stem Cells, 39(2), 156–169. 10.1002/stem.3302

32. Martins, L. R., Sieverling, L., Michelhans, M., Schiller, C., Erkut, C., Grünewald, T. G. P., Triana, S., Fröhling, S., Velten, L., Glimm, H., & Scholl, C. (2024). Single-cell division tracing and transcriptomics reveal cell types and differentiation paths in the regenerating lung. Nature Communications, 15(1), 2246. 10.1038/s41467-024-46469-4

33. MacLeod, A. K., Acosta-Jimenez, L., Coates, P. J., McMahon, M., Carey, F. A., Honda, T., Henderson, C. J., & Wolf, C. R. (2016). Aldo-keto reductases are biomarkers of NRF2 activity and are co-ordinately overexpressed in non-small cell lung cancer. British Journal of Cancer, 115(12), 1530–1539. 10.1038/bjc.2016.363

34. Liberzon, A., Birger, C., Thorvaldsdóttir, H., Ghandi, M., Mesirov, J. P., & Tamayo, P. (2015). The Molecular Signatures Database Hallmark Gene Set Collection. Cell Systems, 1(6), 417–425. 10.1016/j.cels.2015.12.004

35. Dang, C. V. (2012). MYC on the Path to Cancer. Cell, 149(1), 22–35. 10.1016/j.cell.2012.03.003

36. Dongre, A., & Weinberg, R. A. (2019). New insights into the mechanisms of epithelial–mesenchymal transition and implications for cancer. Nature Reviews Molecular Cell Biology, 20(2), 69–84. 10.1038/s41580-018-0080-4

37. Sorensen, G. L. (2018). Surfactant Protein D in Respiratory and Non-Respiratory Diseases. Frontiers in Medicine, 5, 18. 10.3389/fmed.2018.00018

38. Yoneda, M., Xu, L., Kajiyama, H., Kawabe, S., Paiz, J., Ward, J. M., & Kimura, S. (2016). Secretoglobin Superfamily Protein SCGB3A2 Alleviates House Dust Mite-Induced Allergic Airway Inflammation in Mice. International Archives of Allergy and Immunology, 171(1), 36–44. 10.1159/000450788

39. Zeng, Y., Chen, H., Zhang, Z., Fan, J., Li, J., Zhou, S., Wang, N., Yan, S., Cao, J., Liu, J., Zhou, Z., & Liu, W. (2023). IFI44L as a novel epigenetic silencing tumor suppressor promotes apoptosis through JAK/STAT1 pathway during lung carcinogenesis. Environmental Pollution, 319, 120943. 10.1016/j.envpol.2022.120943

40. Eckert, R. L., Fisher, M. L., Grun, D., Adhikary, G., Xu, W., & Kerr, C. (2015). Transglutaminase is a tumor cell and cancer stem cell survival factor. Molecular Carcinogenesis, 54(10), 947–958. 10.1002/mc.22375

41. Lal-Nag, M., Battis, M., Santin, A. D., & Morin, P. J. (2012). Claudin-6: A novel receptor for CPE-mediated cytotoxicity in ovarian cancer. Oncogenesis, 1(11), e33–e33. 10.1038/oncsis.2012.32

42. Li, Y., Lan, T., Liu, M., Li, C., & Du, Y. (2025). GTSF1 promotes stemness in uterine carcinosarcoma through CCL1-mediated M1 macrophage aggregation. American Journal of Cancer Research, 15(5), 2397–2412. 10.62347/MAXH5742

43. Chen, H., Matsumoto, K., Brockway, B. L., Rackley, C. R., Liang, J., Lee, J.-H., Jiang, D., Noble, P. W., Randell, S. H., Kim, C. F., & Stripp, B. R. (2012). Airway Epithelial Progenitors Are Region Specific and Show Differential Responses to Bleomycin-Induced Lung Injury. Stem Cells, 30(9), 1948–1960. 10.1002/stem.1150

44. Nakayama, I., Qi, C., Chen, Y., Nakamura, Y., Shen, L., & Shitara, K. (2024). Claudin 18.2 as a novel therapeutic target. Nature Reviews Clinical Oncology, 21(5), 354–369. 10.1038/s41571-024-00874-2

45. Thijssen, V. L. J. L., Postel, R., Brandwijk, R. J. M. G. E., Dings, R. P. M., Nesmelova, I., Satijn, S., Verhofstad, N., Nakabeppu, Y., Baum, L. G., Bakkers, J., Mayo, K. H., Poirier, F., & Griffioen, A. W. (2006). Galectin-1 is essential in tumor angiogenesis and is a target for antiangiogenesis therapy. Proceedings of the National Academy of Sciences, 103(43), 15975–15980. 10.1073/pnas.0603883103

46. Sidenius, N., & Blasi, F. (2003). The urokinase plasminogen activator system in cancer: Recent advances and implication for prognosis and therapy. Cancer and Metastasis Reviews, 22(2–3), 205–222. 10.1023/A:1023099415940

47. Duch, P., Díaz-Valdivia, N., Ikemori, R., Gabasa, M., Radisky, E. S., Arshakyan, M., Gea-Sorlí, S., Mateu-Bosch, A., Bragado, P., Carrasco, J. L., Mori, H., Ramírez, J., Teixidó, C., Reguart, N., Fillat, C., Radisky, D. C., & Alcaraz, J. (2022). Aberrant TIMP-1 overexpression in tumor-associated fibroblasts drives tumor progression through CD63 in lung adenocarcinoma. Matrix Biology, 111, 207–225. 10.1016/j.matbio.2022.06.009

48. Huang, W., Chiquet-Ehrismann, R., Moyano, J. V., Garcia-Pardo, A., & Orend, G. (2001). Interference of tenascin-C with syndecan-4 binding to fibronectin blocks cell adhesion and stimulates tumor cell proliferation. Cancer Research, 61(23), 8586–8594.

49. Wei, J., Yu, W., Wu, L., Chen, Z., Huang, G., Hu, M., & Du, H. (2023). Intercellular Molecular Crosstalk Networks within Invasive and Immunosuppressive Tumor Microenvironment Subtypes Associated with Clinical Outcomes in Four Cancer Types. Biomedicines, 11(11), 3057. 10.3390/biomedicines11113057

50. Leitinger, B. (2014). Discoidin Domain Receptor Functions in Physiological and Pathological Conditions. In International Review of Cell and Molecular Biology (Vol. 310, pp. 39–87). Elsevier. 10.1016/B978-0-12-800180-6.00002-5

51. Ye, X., Wang, Y., Cahill, H., Yu, M., Badea, T. C., Smallwood, P. M., Peachey, N. S., & Nathans, J. (2009). Norrin, Frizzled-4, and Lrp5 Signaling in Endothelial Cells Controls a Genetic Program for Retinal Vascularization. Cell, 139(2), 285–298. 10.1016/j.cell.2009.07.047

52. Beauchemin, N., & Arabzadeh, A. (2013). Carcinoembryonic antigen-related cell adhesion molecules (CEACAMs) in cancer progression and metastasis. Cancer and Metastasis Reviews, 32(3–4), 643–671. 10.1007/s10555-013-9444-6

53. Zheng, M.-Y., Lin, Y.-T., Kuo, Y.-S., Lin, Y.-J., Kuo, M.-H., Huang, T.-W., Shieh, Y.-S., Huang, Y., & Chou, Y.-T. (2025). Cytokine and epigenetic regulation of CEACAM6 mediates EGFR-driven signaling and drug response in lung adenocarcinoma. Npj Precision Oncology, 9(1), 115. 10.1038/s41698-025-00910-z

54. Hommel, T., Meisel, P. F., Camera, E., Bottillo, G., Teufelberger, A. R., Benezeder, T. H., Wolf, P., Kleissl, L., Stary, G., Posch, C., Schneider, M. R., & Dahlhoff, M. (2025). Loss of ERBB2 and ERBB3 Receptors Impacts Epidermal Differentiation in Mice. Journal of Investigative Dermatology, 145(1), 204–208.e6. 10.1016/j.jid.2024.06.1278

55. Deng, Z., Fan, T., Xiao, C., Tian, H., Zheng, Y., Li, C., & He, J. (2024). TGF-β signaling in health, disease and therapeutics. Signal Transduction and Targeted Therapy, 9(1), 61. 10.1038/s41392-024-01764-w

56. Dcona, M. M., Morris, B. L., Ellis, K. C., & Grossman, S. R. (2017). CtBP- an emerging oncogene and novel small molecule drug target: Advances in the understanding of its oncogenic action and identification of therapeutic inhibitors. Cancer Biology & Therapy, 18(6), 379–391. 10.1080/15384047.2017.1323586

57. 57. Jing, J., Liu, J., Wang, Y., Zhang, M., Yang, L., Shi, F., Liu, P., & She, J. (2018). The role of ZBTB38 in promoting migration and invasive growth of bladder cancer cells. Oncology Reports. 10.3892/or.2018.6937

58. Wang, Y., Ma, C., Yang, X., Gao, J., & Sun, Z. (2024). ZNF217: An Oncogenic Transcription Factor and Potential Therapeutic Target for Multiple Human Cancers. Cancer Management and Research, Volume 16, 49–62. 10.2147/CMAR.S431135

59. Ingram, K., Samson, S. C., Zewdu, R., Zitnay, R. G., Snyder, E. L., & Mendoza, M. C. (2022). NKX2-1 controls lung cancer progression by inducing DUSP6 to dampen ERK activity. Oncogene, 41(2), 293–300. 10.1038/s41388-021-02076-x

60. Lin, Y., Lin, P., Guo, W., Huang, J., Xu, X., & Zhuang, X. (2023). PLAGL2 promotes the stemness and is upregulated by transcription factor E2F1 in human lung cancer. Environmental Toxicology, 38(4), 941–949. 10.1002/tox.23739

61. Zhu, J., Wu, C., Li, H., Yuan, Y., Wang, X., Zhao, T., & Xu, J. (2016). DACH1 inhibits the proliferation and invasion of lung adenocarcinoma through the downregulation of peroxiredoxin 3. Tumor Biology, 37(7), 9781–9788. 10.1007/s13277-016-4811-x

62. Rockich, B. E., Hrycaj, S. M., Shih, H. P., Nagy, M. S., Ferguson, M. A. H., Kopp, J. L., Sander, M., Wellik, D. M., & Spence, J. R. (2013). Sox9 plays multiple roles in the lung epithelium during branching morphogenesis. Proceedings of the National Academy of Sciences, 110(47). 10.1073/pnas.1311847110

63. Kobayashi, W., & Ozawa, M. (2013). The transcription factor LEF-1 induces an epithelial–mesenchymal transition in MDCK cells independent of β-catenin. Biochemical and Biophysical Research Communications, 442(1–2), 133–138. 10.1016/j.bbrc.2013.11.031

64. Zhang, P., Sun, Y., & Ma, L. (2015). ZEB1: At the crossroads of epithelial-mesenchymal transition, metastasis and therapy resistance. Cell Cycle, 14(4), 481–487. 10.1080/15384101.2015.1006048

65. Fu, T., Dai, L.-J., Wu, S.-Y., Xiao, Y., Ma, D., Jiang, Y.-Z., & Shao, Z.-M. (2021). Spatial architecture of the immune microenvironment orchestrates tumor immunity and therapeutic response. Journal of Hematology & Oncology, 14(1), 98. 10.1186/s13045-021-01103-4

66. Coënon, L., Geindreau, M., Ghiringhelli, F., Villalba, M., & Bruchard, M. (2024). Natural Killer cells at the frontline in the fight against cancer. Cell Death & Disease, 15(8), 614. 10.1038/s41419-024-06976-0

67. Gaggero, S., Witt, K., Carlsten, M., & Mitra, S. (2020). Cytokines Orchestrating the Natural Killer-Myeloid Cell Crosstalk in the Tumor Microenvironment: Implications for Natural Killer Cell-Based Cancer Immunotherapy. Frontiers in Immunology, 11, 621225. 10.3389/fimmu.2020.621225

68. Cao, J., Chow, L., & Dow, S. (2023). Strategies to overcome myeloid cell induced immune suppression in the tumor microenvironment. Frontiers in Oncology, 13, 1116016. 10.3389/fonc.2023.1116016

69. Chen, H., Xu, Z., & Varner, J. (2025). Targeting myeloid cells to improve cancer immune therapy. Frontiers in Immunology, 16, 1623436. 10.3389/fimmu.2025.1623436

70. Kim, N., Kim, H. K., Lee, K., Hong, Y., Cho, J. H., Choi, J. W., Lee, J.-I., Suh, Y.-L., Ku, B. M., Eum, H. H., Choi, S., Choi, Y.-L., Joung, J.-G., Park, W.-Y., Jung, H. A., Sun, J.-M., Lee, S.-H., Ahn, J. S., Park, K., … Lee, H.-O. (2020). Single-cell RNA sequencing demonstrates the molecular and cellular reprogramming of metastatic lung adenocarcinoma. Nature Communications, 11(1), 2285. 10.1038/s41467-020-16164-1

71. Laughney, A. M., Hu, J., Campbell, N. R., Bakhoum, S. F., Setty, M., Lavallée, V.-P., Xie, Y., Masilionis, I., Carr, A. J., Kottapalli, S., Allaj, V., Mattar, M., Rekhtman, N., Xavier, J. B., Mazutis, L., Poirier, J. T., Rudin, C. M., Pe’er, D., & Massagué, J. (2020). Regenerative lineages and immune-mediated pruning in lung cancer metastasis. Nature Medicine, 26(2), 259–269. 10.1038/s41591-019-0750-6

72. Han, G., Sinjab, A., Rahal, Z., Lynch, A. M., Treekitkarnmongkol, W., Liu, Y., Serrano, A. G., Feng, J., Liang, K., Khan, K., Lu, W., Hernandez, S. D., Liu, Y., Cao, X., Dai, E., Pei, G., Hu, J., Abaya, C., Gomez-Bolanos, L. I., … Kadara, H. (2024). An atlas of epithelial cell states and plasticity in lung adenocarcinoma. Nature, 627(8004), 656–663. 10.1038/s41586-024-07113-9

73. Sikkema, L., Ramírez-Suástegui, C., Strobl, D. C., Gillett, T. E., Zappia, L., Madissoon, E., Markov, N. S., Zaragosi, L.-E., Ji, Y., Ansari, M., Arguel, M.-J., Apperloo, L., Banchero, M., Bécavin, C., Berg, M., Chichelnitskiy, E., Chung, M., Collin, A., Gay, A. C. A., … Theis, F. J. (2023). An integrated cell atlas of the lung in health and disease. Nature Medicine, 29(6), 1563–1577. 10.1038/s41591-023-02327-2

